# Characterization of an ECF56-family sigma factor from *Streptomyces venezuelae* reveals a highly conserved regulome

**DOI:** 10.1101/2020.01.30.927608

**Authors:** Mitchell G. Thompson, Amin Zargar, Pablo Cruz-Morales, Tristan De Rond, Samantha Chang, Allison N. Pearson, Garima Goyal, Jesus F. Barajas, Jacquelyn M. Blake-Hedges, Ryan M. Phelan, Victor Reyes-Umana, Amanda C. Hernández, Nathan J. Hillson, Patrick M. Shih, Jay D. Keasling

## Abstract

Bacteria often possess alternative sigma factors that initiate the transcription of specific genes under environmental stresses, the largest and most diverse group being the extracytoplasmic function (ECF) sigma factors. The regulation of ECF activity is crucial for ensuring the distinct transcription of stress responsive genes only occurs under the appropriate conditions. While most ECFs are comprised of only the core σ_2_ and σ_4_ regions, a unique form of ECF sigma factor regulation also contains a C-terminal extension bearing homology to the NTF2 superfamily of protein domains. While previous work has shown that this NTF2 domain can affect transcriptional activity *in vivo* in ECF41 and ECF42, its role in the newly classified ECF56 subgroup is unknown. In this work, we show that truncation of the C-terminus of the ECF56 sigma factor SVEN_4562 of *Streptomyces venezuelae* upregulates its activity in a hybrid assay. Through transcriptomics in *S. venezuelae*, we found that this truncated ECF56 sigma factor has a highly conserved promoter sequence *in vivo*. Bioinformatic assays illustrated that deep branches of the *Actinobacteria* phylum contained putative ECF56 promoter motifs identical to those found in the *S. venezuelae* ECF56 regulon. We validated these findings through *ex situ* hybrid assays illustrating that truncated ECF56 sigma factors from phylogenetically diverse *Actinobacteria* activate transcription from these promoters. Importantly, our work shows that the genetic infrastructure of the ECF56 family of sigma factors is highly conserved and performs important functions yet to be understood in *Actinobacteria*.

**Importance:** Most ECF sigma-factors rely on anti-sigma factor regulation; in contrast, the unique classes of ECF sigma-factors that contain a C-terminal extension are thought to respond directly to an environmental signal. Here we show that the *cis-*acting regulatory element of the ECF56 regulon is likely highly conserved in many *Actinobacteria*, with exact nucleotide level conservation over ~2 billion years of evolution. The high conservation of this genetic architecture, as well as a conserved gene content within the regulon, strongly point to a specialized and important role in *Actinobacteria* biology.

## Introduction

Sigma factors are required for the initiation of transcription in all known bacteria. They are able to recruit the RNA polymerase (RNAP) core, direct it to a specific promoter region, and initiate transcription by enabling promoter melting (1). While most genes in a given bacterium are transcribed using a “housekeeping” sigma factor, bacteria often possess alternative sigma factors that initiate transcription of specific subsets of genes in response to specific environmental stresses (1). Families of alternative sigma factors are involved in responses to stress such as responding to heat shock (RpoH-like), flagellar synthesis (FliA-like), and nitrogen limitation (RpoN-like) (1). The largest and most diverse group of alternative sigma factors are the Extracytoplasmic Function (ECF) sigma factors (1). ECF sigma factors are smaller in size compared to housekeeping σ^70^ sigma factors, and normally possess only two of the four conserved domains, σ_2_ and σ_4_. These domains are sufficient to recognize the −10 and −35 elements of a bacterial promoter, recruit RNAP core, and initiate transcription via promoter melting (1). Despite their reduced size, ECF sigma factors play a role of outsized importance in the transduction of environmental signals in bacteria (2–4).

The regulation of ECF activity is critical to ensure that the transcription of stress responsive genes only occurs under the appropriate conditions. In many cases, ECFs are negatively regulated by cognate anti-sigma factors, which are usually co-transcribed as part of a negative feedback loop (5). Anti-sigma factors physically sequester ECFs until an appropriate environmental signal either leads to the physical destruction of the anti-sigma factor or alters the affinity of the anti-sigma factor for its cognate ECF (6). The mechanism that causes dissociation varies widely depending on the ECF sigma factor, but can take the form of a proteolysis event, export of the anti-sigma factor, or inactivation of the anti-sigma factor through phosphorylation (5–7).

Another form of ECF sigma factor regulation has been described more recently, via C-terminal extensions of ~100 amino acids that show homology to members of the NTF2 superfamily of protein domains (8). The NTF2 superfamily of domains is widely distributed on the tree of life and shows little conservation at the primary sequence identity level; but members include many enzymes and a recent review of their function has implicated allosteric regulation of DNA binding proteins (8). Studies of these NTF2 domains in ECF41 and ECF42 found it was critical to initiation of gene transcription and strongly implicate that it directly responds to an external signal (9–12). In 2015, a genomic survey of ECFs and other signal transduction proteins in *Actinobacteria* revealed another group of ECFs with similar C-terminal extensions: the ECF56 subgroup (13). Previous work with the *Mycobacterium tuberculosis* ECF56 family sigma factor Rv0182c (*sigG*) implicated involvement in DNA damage response pathways (14, 15). While genes regulated by SigG were identified, neither the physiological role these genes play nor the environmental cue SigG responds to were identified. To date, the specific signal that is transduced by the NTF2 domain of ECF56 or any other ECF family members has yet to be identified.

In an attempt to shed light on the physiological role of the ECF56 subfamily, we characterized the regulatory activity of the NTF2 domain of ECF56 sigma factors *ex situ* in *Escherichia coli* and *in vivo* in *S. venezuelae*. Beyond its attributes as a fast-growing, genetically tractable *Actinobacteria*, *S. venezuelae* has also been previously used to study the NTF2 domain of the ECF42 sigma factor subfamily (10). In this work, we identify a highly conserved promoter that defines an ECF56 regulon in *S. venezuelae* and demonstrate through bioinformatics that the regulon is highly conserved through deep branches of *Actinobacteria*. Lastly, we further validate our findings regarding the role of these C-terminal extensions by determining their role in diverse ECF56 sigma factors.

## Results

### The C-terminal extension of ECF56 sigma factors inhibits transcription

We first aimed to determine the regulatory role of these C-terminal extensions in ECF56 sigma factors. To start, we sought to determine the differential transcriptional activity with and without the C-terminal extension of the ECF56 sigma factor from *S. venezuelae*, SVEN_4562 (**Figure 1A**). Previously, Wecke and colleagues had identified a conserved DGGGK motif downstream of the σ_4_ domain in ECF41 family sigma factors (**Figure S1A**). Truncating the C-terminus of these sigma factors immediately after this motif resulted in increased transcription from cognate promoters (12). By aligning >900 ECF56 sigma factors, we observed a highly conserved MPP(F/Y/L) motif similarly positioned relative to the σ_4_ domain (**Figure S1B**). As ~60% of ECF56 sigma factor sequences contained an aromatic phenylalanine or tyrosine after the highly conserved MPP, we truncated SVEN_4562 immediately after its internal MPPY to maintain the aromatic residue, resulting in SVEN_4562_Δ237-323_.

**Figure 1.**
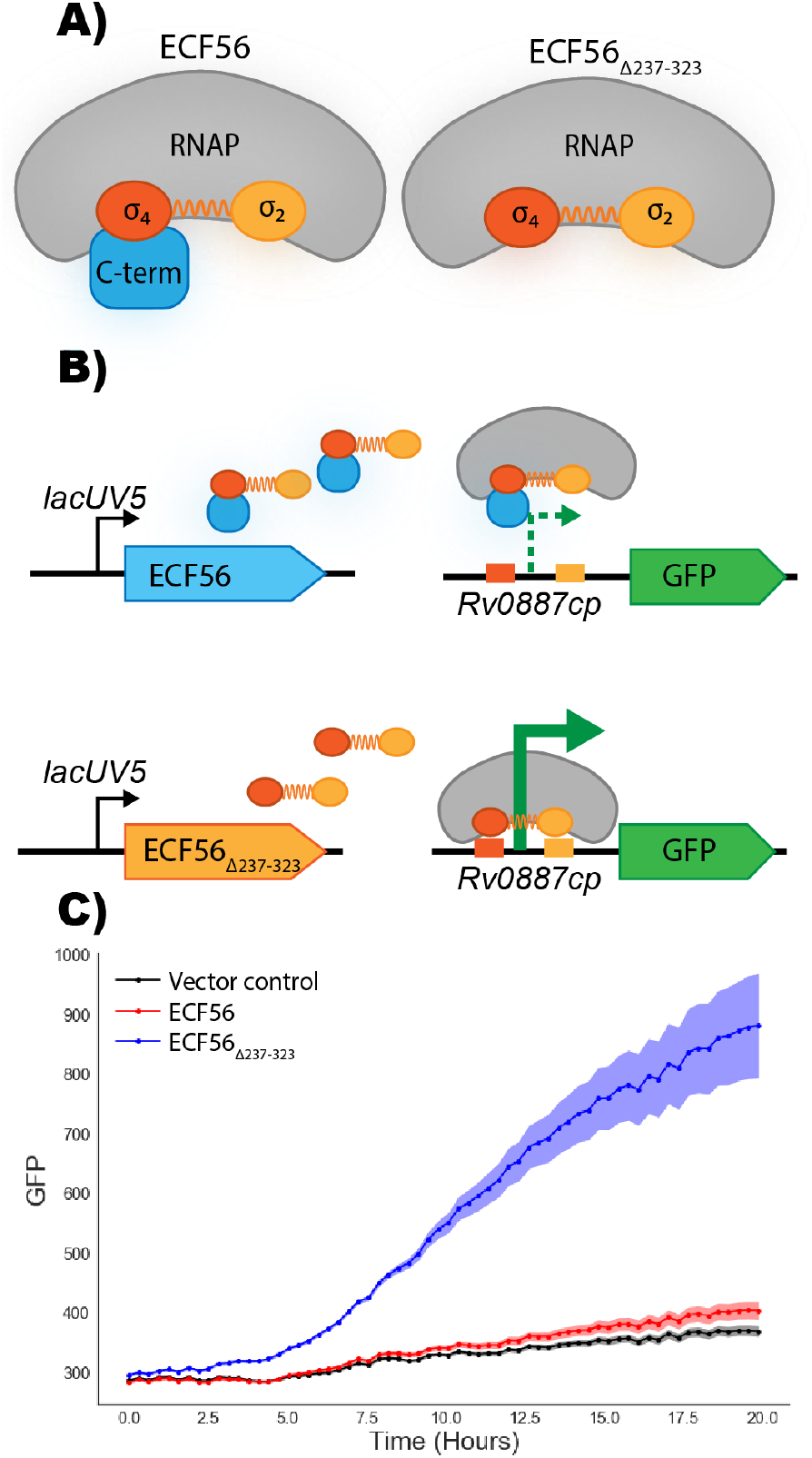
Architecture, and characterization of ECF56 sigma factors. **A)** Schematic illustrates the interactions between RNA polymerase and ECF sigma factors with and without the NTF2 domain. **B)** The schematic illustrates the genetic circuit built in *E. coli*. Wild-type ECF56 sigma factor(SVEN_4562) or a truncated variant is constitutively expressed and results in the transcription of a GFP cassette driven by the known ECF56-dependent promoter. **C)** Graph shows GFP fluorescence in arbitrary units over time in strains expressing variants of SVEN_4562 or an empty vector control. Shaded area shows 95% confidence intervals, n=3.

To test this hypothesis we measured transcription from a previously identified *Rv0182c* target promoter (*Rv0887cp*) from *M. tuberculosis* in an *ex situ* assay (15). In *E. coli*, the transcription of each sigma factor was driven by a leaky uninduced lacUV5 promoter, and its ability to bind and drive expression of the *Rv0887c* promoter was determined through a GFP reporter protein (**Figure 1B**). Compared to the empty vector control, we observed a 2.4-fold increase in GFP expression in the strain overexpressing the truncated ECF56 sigma factor. In contrast, overexpression of the full ECF56 sigma factor resulted in a 1.1-fold increase in GFP expression. These observations suggest that the C-terminal extension of ECF56 sigma factors inhibits promoter binding. Interestingly, this contrasts with other investigations of this domain in *Actinobacteria*, which found that the full C-terminal extension was required for complete activation of transcription (10–12).

### Defining the ECF56 Regulon of *S. venezuelae*

As the ECF56 family is widely distributed in *Actinobacteria (13)*, we hypothesized that they must play an important physiological role in these organisms, such as *S. venezuelae*. Therefore, we cloned the truncated SVEN_4562 under the strong constitutive promoter kasOp and genetically inserted it into the S. venezuelae genome at the *phiC* attachment site. Constitutive expression of truncated SVEN_4562 produced a bald phenotype (**Figure 2A**). This is especially interesting because C-terminal truncations of ECF42 in *S. venezuelae* showed no observable differences in morphology (10).

**Figure 2.**
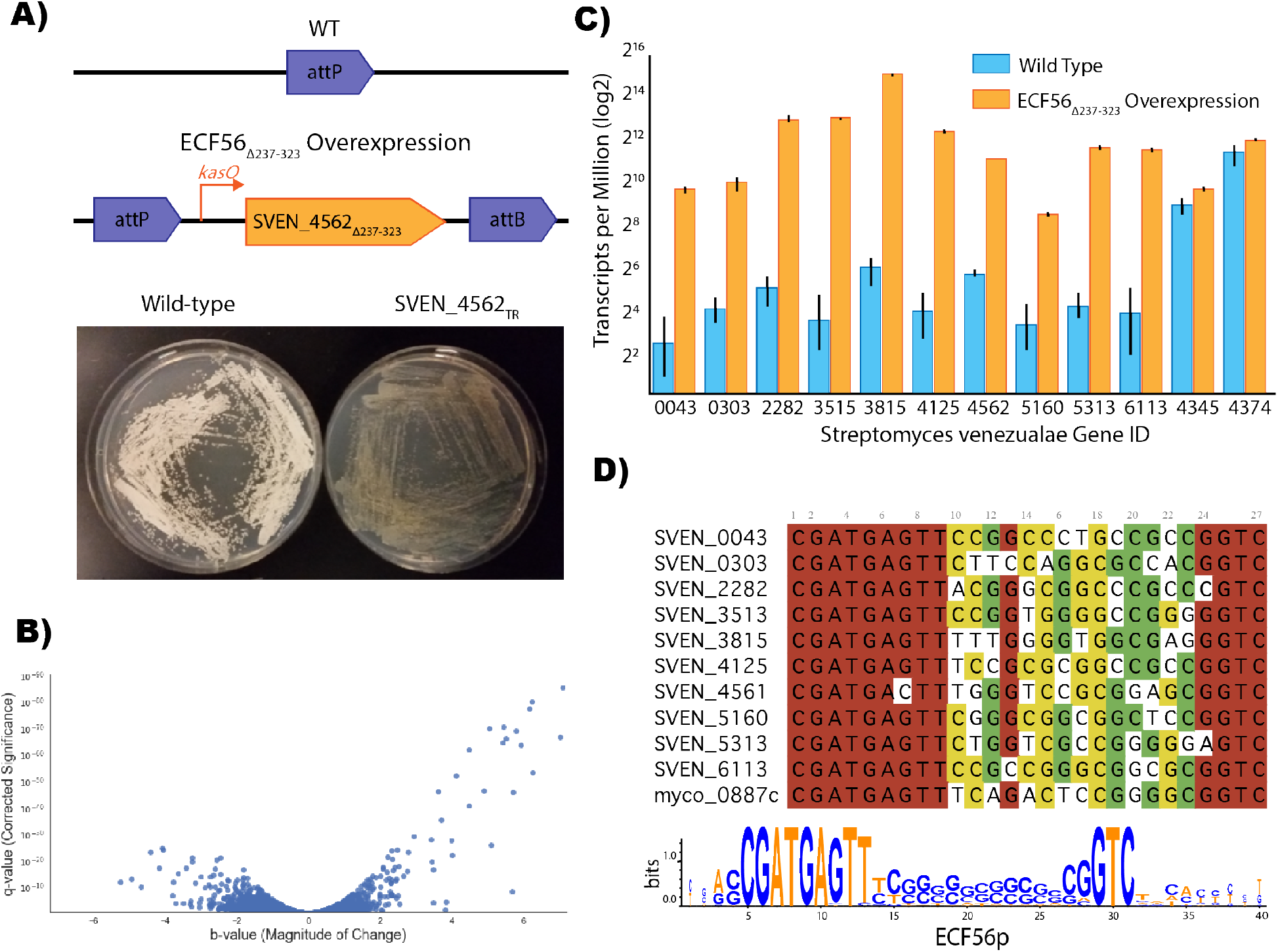
Identification of genes regulated by SVEN_4562. **A)** A truncated version of ECF56 from *S. venezuelae* (SVEN_4562) driven by the kasOp promoter was inserted into the genome of *S. venezuelae* resulting in a bald phenotype when compared to the wild-type control. **B)** Volcano plot showing RNA-Seq analysis of transcripts of the truncated SVEN_4562 overexpressing strain compared to wild-type *S. venezuelae*. The y-axis shows the q-value (corrected significance), while the x-axis shows b-value (magnitude of change). Results represent n=3 **C)** Genes believed to be directly upregulated when truncated SVEN_4562 is overexpressed, in addition to *rpoD* (SVEN_4345) and *EF-Tu* (SVEN_4374). Error bars show 95% confidence interval, n=3. **D)** Top panel illustrates an alignment of the upstream sequences of these upregulated genes and the bottom panel represents a position weight matrix of these sequences.

Through transcriptomics, we sought to develop a greater understanding of the physiological role of ECF56 in S. venezuelae. Using RNA sequencing (RNA-seq), we compared the transcriptomes of a strain overexpressing the truncated ECF56 to the wild-type strain and observed a population of genes that were heavily upregulated (**Figure 2B**). The complete list of differentially expressed genes can be found as **Supplemental Attachment 1**. Our results showed that 22 loci had a *b* of >3.5 and a *q-value* less than 1×10^-10^, representing the most highly upregulated genes. However, previous work in studies where transcription factors were overexpressed had noticed significant read through between genes that could potentially confound results (16). To account for this possibility we manually investigated reads mapped to the *S. venezuelae* chromosome to identify instances in which increased differential expression was likely due to read through (or direct overexpression in the case of SVEN_4562) and identified 12 genes likely to be false hits (**Table S1**). The resulting genes that were specifically upregulated by the truncated ECF56 sigma factor were amongst the most highly expressed in the cell, with transcript per million (TPM) values either close to or as high as genes such as *rpoD* and *EF-Tu* **(Figure 2C)**. Unfortunately, none of the genes found to be upregulated and controlled by SVEN_4562 directly had a known function. Of the ten genes, six were annotated as proteins of hypothetical function, whereas the remaining genes were annotated in broadly functional families such as glyoxalase, hydrolase, dihydrofolate reductase, and deaminase-reductase (**Table 1**). Interrogation of the upstream sequences of the genes in **Table 1** revealed a nearly perfectly conserved promoter motif (CGATGAGTTN_14_GGTC - **Figure 2D**). Even more striking was that this promoter motif was also perfectly conserved in the distantly related *M. tuberculosis* ECF56 sigma factor regulated *Rv0887c* (**Figure 2D**).

**Table 1:** Genes predicted to be regulated by SVEN_4562. Genes that were found to be highly upregulated during the overexpression of truncated SVEN_4562, and not to be the result of suspected readthrough. The b-value is a measure of change in expression (17), where as the qval is a corrected significance value.

| Gene ID | Description | *b* | *q-value* |
| --- | --- | --- | --- |
| SVEN_3513 | putative hydrolase | 7.08 | 3.89E-86 |
| SVEN_3815 | hypothetical protein | 6.52 | 2.47E-93 |
| SVEN_4125 | hypothetical protein | 6.24 | 8.38E-81 |
| SVEN_6113 | Dihydrofolate reductase | 5.92 | 2.27E-64 |
| SVEN_2282 | Glyoxalase family protein | 5.79 | 8.97E-70 |
| SVEN_0043 | hypothetical protein | 5.71 | 2.58E-46 |
| SVEN_5313 | Bifunctional deaminase-reductase domain protein | 5.50 | 1.41E-66 |
| SVEN_0303 | hypothetical protein | 4.47 | 1.98E-41 |
| SVEN_5160 | hypothetical protein | 4.13 | 1.33E-52 |
| SVEN_4561 | hypothetical protein | 3.70 | 4.14E-36 |

### Predicting the ECF56 Regulome

Given the highly conserved nature of the *S. venezuelae*, and *M. tuberculosis* promoters we sought to investigate the genomic context of ECF56 homologs in selected Actinobacterial genomes. We searched for homologs of ECF56 in a database including 720 high quality genomes from actinobacterial species. We retrieved 599 homologs from 414 genomes, mostly from Streptomycetaceae, Mycobacteria and Pseudonocardiaceae. We found that most *Streptomyces spp.* (75%) included a single ECF56 homolog. A phylogenetic reconstruction (**Supplemental Attachment 2**) showed that these single copy-ECF56 homologs are monophyletic, we also found that these homologs share a high degree of synteny among streptomycetes.

After confirming the broad distribution and conservation of the ECF56 we then sought to identify putative members of regulons in distinct species. For this analysis we selected ten representative genomes and searched for the ECF56 consensus binding sequence “CGATGAGTTN_15_GTC”. This search revealed that many proteins belonging to the glyoxylase, dehydrofolate reductase and alpha/beta hydrolase families among others are under the control of ECF56 across largely divergent actinobacterial species. Two interesting exceptions were found: only one protein (a member of the glyoxylase family) is located upstream of the ECF56 binding sequence in *M tuberculosis* while none could be found in *Salinispora tropica* which nevertheless includes a bonafide ECF56 homolog.

### Conservation of target promoter of ECF56 across phyla

Lastly, we wanted to investigate the potential upregulation of transcription of the truncated ECF56 in various species, as similar studies in ECF41 have shown varying results depending on the coding sequence. As such, we cloned truncated versions of two more ECF56 from *C. woesei* and *M. tuberculosis* (**Figure 4A**). *C. woesei* was selected due to its divergence from Streptomycetes ~2 billion years ago (18), yet bioinformatic analysis revealed the identical promoter sequences were conserved in its genome. *M. tuberculosis* was selected due to its importance as a public health threat, and previous work that has been done on the Rv0182c regulon (14, 15). As with SVEN_4562, both of these homologs were truncated immediately past a conserved MPPF motif in the case of Cwoe_2709, and an MPPY motif for Rv0182c. We also cloned an additional ECF dependent promoter, SVEN_3513p, to compare the differences due to the regions varying between the −35 and −10 elements. All ECF56 truncations performed similarly in the *ex situ* assay, while the promoter of M. tuberculosis showed greater upregulation than the promoter of SVEN_3513, suggesting that the intergenic −35 and −10 region does affect transcription.

## Discussion

Traditionally, sigma factors have been thought to rely on one or more trans-acting partners to transduce specific environmental cues in order to activate responsive gene expression (5). Recent work has challenged this notion, with the discovery that many C-terminal extensions modulate the activity of a variety of ECF sigma factor families (19). These domains presumably are able to sense environmental signals, which results in conformational changes that activate expression of their cognate regulon, effectively turning them into one-component systems (19). However despite recent efforts, which have defined multiple putative regulons of these ECF sub-families, the physiological roles that these ECFs regulate and the signals that they respond to remain a mystery (19).

With its unique role in ECF transcription, efforts have been made to study the NTF2 domain and its interaction with the target promoter and RNA polymerase. Initial studies in the ECF41 subfamily in *B. lincheniformis* (Firmicutes) and *R. sphaeroides* (α-proteobacteria) found that truncations of the NTF2 domain drastically upregulated transcription of a highly conserved promoter (12). Interestingly, biochemical and crystal structure studies of the ECF41 σ^J^ of *M. tuberculosis* (*Actinobacteria*) and the ECF42 of S. venezuelae (*Actinobacteria*) revealed that the NTF2 domain was necessary for transcription (9, 10). While a direct coupling analysis of ECF41 and ECF42 revealed that the NTF2 region bind and regulate the core ECF regions of σ_2_ and σ_4_ responsible for transcription (11), the role of these NTF2 domains in the ECF56 subfamily has not yet been studied. Crystallographic studies of ECF56 homologs may provide crucial information on how NTF2 domains impose negative regulation.

The only previously studied member of the ECF56 subfamily is Rv0182c (SigG) of *M. tuberculosis*. SigG was shown to be the most weakly expressed sigma factor in *M. tuberculosis* when grown in laboratory conditions, potentially making it challenging to study (20). However microarray analysis revealed *sigG* to be one of the most upregulated genes when *M. tuberculosis* was grown during human macrophage infection prompting further research (21). Further work then showed that SigG was upregulated in the presence of the DNA damaging agent mitomycin C, and that it controlled the expression of two proteins that contained glyoxalase domains (14, 15). This connection to the DNA-damage response was recently confirmed when it was found that SigG is regulated by the PafBC system (22). PafBC has recently been shown to be the master transcriptional regulator of the LexA/RecA-independent DNA repair response in *M. tuberculosis (22)*. One of the main functions of this response is to proteolytically degrade RecA, to temporally control the stress response (22). This may suggest that the C-terminus of ECF56 sigma factors respond to specific DNA damage signals to activate a highly conserved stress response (**Figure 5**). Our transcriptomic investigation in *S. venezuelae* revealed putative glyoxalase proteins strongly upregulated with constitutive expression of the truncated ECF56 sigma factor. As our bioinformatic studies revealed many other *Actinobacteria* also had glyoxalases, dihydrofolate reductases, and α/β hydrolases in their predicted ECF56 regulons, it is conceivable that these enzymes are required to metabolize toxic byproducts of DNA damage. However, these protein families are known to have a wide range of functions, and further experimentation is clearly required to predict their exact function. We believe that future work should focus heavily on the physiological and biochemical roles these proteins play, which will further elucidate the signals that activate ECF56 sigma factors.

Rhodius et al. had previously proposed that ECF sigma factors may make excellent parts for synthetic biology by demonstrating their exceptional orthogonality (23). They further went on to show that domains from different ECF subfamilies can be combined to generate functional proteins that recognize hybrid promoters (23). From this work various groups have used ECF sigma factors to build highly insulated genetic circuits (24), ECF sigma factor based toolkits for both *E. coli* and *Bacillus subtilis* (25, 26), and highly complex synthetic timers (27). All of these synthetic circuits take advantage of the highly specific interaction ECF sigma factors have with their cognate promoters. Based on the extreme conservation of the ECF56 promoter, the ECF56 subfamily is also likely to be highly orthogonal to other ECF promoters. While this orthogonality is highly desirable in genetic engineering, the activity of most ECF sigma factors relies on control by either cognate anti-sigma factors or synthetic genetic circuitry. This excess genetic baggage may complicate some engineering tasks. The prospect that C-terminal extended ECF sigma factor may be able to transduce a variety of environmental cues is extremely intriguing. If the signal could be identified, these proteins could function as highly orthogonal, tunable, stand alone genetic parts. Further work on these unique transcription factors should heavily focus on deorphaning these receptors.

## Methods

### Media, Chemicals, and Strains

*E. coli* bacterial cultures were grown in Luria-Bertani (LB) Miller medium (BD Biosciences, USA) at 37 °C. *S. venezuelae* ATCC 10712 was grown on MYM media (28), and sporulated on mannitol soy (MS) agar media supplemented with 10 mM magnesium chloride (28). Cultures were supplemented with carbenicillin (100 mg/L, Sigma Aldrich, USA), kanamycin (50 mg/L, Sigma Aldrich, USA), or spectinomycin (100 mg/L for *E. coli* or 400 mg/L for *S. venezuelae*, Sigma Aldrich, USA) All bacterial strains and plasmids used in this work are listed in **Table 2** and are available through the public instance of the JBEI registry. (https://public-registry.jbei.org/).

**Table 2:** Strains and Plasmids used in this study.

| Strains | JBEI Number | Source | Notes |
| --- | --- | --- | --- |
| *E. coli* DH10B |  | Thermo Fisher |  |
| *E. coli* ET12567 |  | ATCC BAA-525 |  |
| *S. venezuelae* |  | ATCC 10712 |  |
| Plasmids |  |  |  |
| pUZ8002 |  | (29) |  |
| pSC-KasOp-mCherry |  | (30) | Spectinomycin |
| pSC-KasOp-SVEN_4562_trunc | TBD | This work | Spectinomycin |
| pBbE5k-RFP |  | (31) | Kan |
| pBbB5a-RFP |  | (31) | Amp |
| pBbE5k-SVEN_4562 | TBD | This work | Kan |
| pBbE5k-SVEN_4562_trunc | TBD | This work | Kan |
| pBbE5k-Cwoe_2709 | TBD | This work | Kan |
| pBbE5k-Cwoe_2709_trunc | TBD | This work | Kan |
| pBbE5k-Rv0182c | TBD | This work | Kan |
| pBbE5k-Rv0182c_trunc | TBD | This work | Kan |
| pBbBa Rv0877cp::GFP | TBD | This work | Amp |
| pBbBa SVEN_3513p::GFP | TBD | This work | Amp |

### DNA Manipulation

All plasmids were designed using Device Editor and Vector Editor software, while all primers used for the construction of plasmids were designed using j5 software (32–34). Plasmids were assembled via Gibson Assembly using standard protocols (35), or Golden Gate Assembly using standard protocols(36). Plasmids were routinely isolated using the Qiaprep Spin Miniprep kit (Qiagen, USA), and all primers were purchased from Integrated DNA Technologies (IDT, Coralville, IA). *E. coli* codon optimized gene blocks for *Rv0182c* and Cwoe_2709 were purchased from Integrated DNA Technologies (IDT, Coralville, IA).

### Conjugation into *Streptomyces venezuelae*

*E. coli* ET12567/pUZ8002 was transformed with an appropriate vector and selected for on LB agar containing kanamycin (25 μg/mL), chloramphenicol (15 μg/mL), and apramycin (50 μg/mL). A single colony was used to inoculate 5 mL of LB containing kanamycin (25 μg/mL), chloramphenicol (15 μg/mL), and apramycin (50 μg/mL) at 37°C. The overnight culture was used to seed 10 mL of LB containing the same antibiotics, which was grown at 37°C to an OD600 of 0.4-0.6. The *E. coli* cells were pelleted by centrifugation, washed twice with LB, and resuspended in 500 μL of LB. Fresh *S. venezuelae* J1074 spores were collected from a mannitol soy agar plate with 5 mL of 2xYT and incubated at 50°C for 10 min. The spores (500 μL) and the *E. coli* cells (500 μL) were mixed, spread onto mannitol soy agar, and incubated at 30°C for 16 hours. After 1 mL addition of nalidixic acid (20 μg/mL) and apramycin (40 μg/mL) was added and allowed to dry, the plate was further incubated for 3-4 days at 30°C. A single colony was used to inoculate into TSB containing nalidixic acid (25 μg/mL) and apramycin (25 μg/mL). The overnight culture was spread onto a MS plate and incubated at 30°C for 2-3 days. The spores were collected from the plate with 3-4 mL of water and mixed with glycerol to prepare 25% glycerol stock. The glycerol stock was stored at - 80°C for long-term storage.

### RNA Isolation and RNA-Seq

RNA was isolated from mycelial cultures grown for 24 hours on MYM media at 30 °C. Briefly, 1 mL of culture was mixed with 2 volumes of RNAlater (Invitrogen, USA) for five minutes at room temperature, pelleted in a Eppendorf 5245 centrifuge for 2 minutes at 10,000 rpm and decanted. Pelleted mycelia were then stored at −80 °C until RNA was extracted. RNA was isolated using QIAGEN RNeasy kit (Qiagen, USA) per manufacturer’s instructions for gram-positive bacteria. Genomic DNA was removed with Ambion Turbo DNase (Invitrogen, USA). Bacterial rRNA was removed from total RNA using a Ribo-Zero RNA removal kit (Illumina, USA), and libraries for RNAseq were prepared using the NEBNext Ultra II RNA Library Prep Kit for Illumina (New England Biolabs, USA) according to the manufacturer’s instructions. Libraries were then sequenced on a Illumina MiSeq using kit v3 (2 × 75-cycle) with paired-end reads. RNAseq data was analyzed with Kallisto (37) which pseudo-aligned reads to the *S. venezuelae* ATCC 10712 coding sequences (GCA_008639165.1). Sleuth was then used to calculate differential expression (17). Raw transcript per million (TPM) values, genome wide differential expression, and the top 100 most upregulated genes during SVEN_4562 _trunc overexpression can be found in **Supplemental Attachment 1**. In order to identify potential promoter readthrough, raw reads were mapped to the *S. venezuelae* ATCC 10712 genome using Bowtie 2 (38), and mapped reads were visualized with the Integrated Genomics Viewer (39). Genes that showed upregulation as a consequence from readthrough were not considered for further promoter analysis.

### Fluorescence Assays

Fluorescence assays were conducted as previously described with minor changes (40). All endpoint assays were conducted in 96-deep well plates (Corning Costar, 3960), with each well containing 500 **μ**L of LB medium with appropriate antibiotics inoculated at 1% v/v from overnight cultures. Plates were sealed with AeraSeal film (Excel Scientific, AC1201-02 and incubated at 37 C in a 250 rpm shaker rack. After 24 hours, 100 **μ**L from each well was aliquoted into a black, clear-bottom 96-well plate for measurements of optical density and fluorescence using an Infinite F200Pro (Tecan Life Sciences, San Jose, CA) plate reader. Optical density was measured at 600 nm (OD_600_), while fluorescence was measured using an excitation wavelength of 470 nm, an emission wavelength of 530 nm, and a manually set gain of 25. To perform time course assays, overnight cultures were inoculated into 10 mL of LB medium from single colonies, and were grown at 37 °C. These cultures were then diluted 1:100 into 100 uL of LB medium in 96-well plates (Falcon, 353072). Plates were sealed with a gas-permeable microplate adhesive film (VWR, USA), and then optical density and fluorescence were monitored for 20 hours in a Tecan Infinite F200Pro (Tecan Life Sciences, San Jose, CA) plate reader. Optical density was measured at 600 nm (OD_600_), while fluorescence was measured using an excitation wavelength of 470 nm, an emission wavelength of 530 nm, and a manually set gain of 25.

### Bioinformatics

Sequences of ECF sigma factors were downloaded from Pfam - https://pfam.xfam.org/family/Sigma70_ECF. An ECF56 sigma factor HMM was generated as described previously (4), and ECF56 family sigma factors were identified using hmmscan (41). ECF56 alignments were done using the MAFFT-LINSI algorithm (42) and visually inspected using UGENE software (43). Alignments were restricted to positions that had greater than 50% occupancy using Protein Dynamics and Sequence Analysis (ProDy) (44, 45). All conserved sequence motifs were visualized with Weblogo (46).

We mined for ECF56 homologs in actinobacteria using the amino acid sequence of SVEN_4562 as query for searches using BlastP with an E-value cutoff of1E-12 and a bit-score cutoff of 200. Independent BlastP searches were done with each one of 720 high quality genomes. The retrieved protein sequences were aligned using muscle (47), and trimmed using jalview (48). An ECF56 phylogenetic tree was obtained using IQtree with the resulting amino acid matrix the best amino acid substitution model for the tree was selected using the ModelFinder utility implemented in IQtree (49). Node support comes from 10000 bootstrap generations. For analysis of the genome context of the selected ECF56 homologs shown in figure 3 we used CORASON with an adhoc genomes database including selected organisms (50). the parameters for the search were e-value 1E-12 and bit-score cutoff of 200, the graphical output was modified to follow the order of a phylogenetic tree generated with the protein sequences homologs identified by CORASON with identical parameters to those described above.

**Figure 3.**
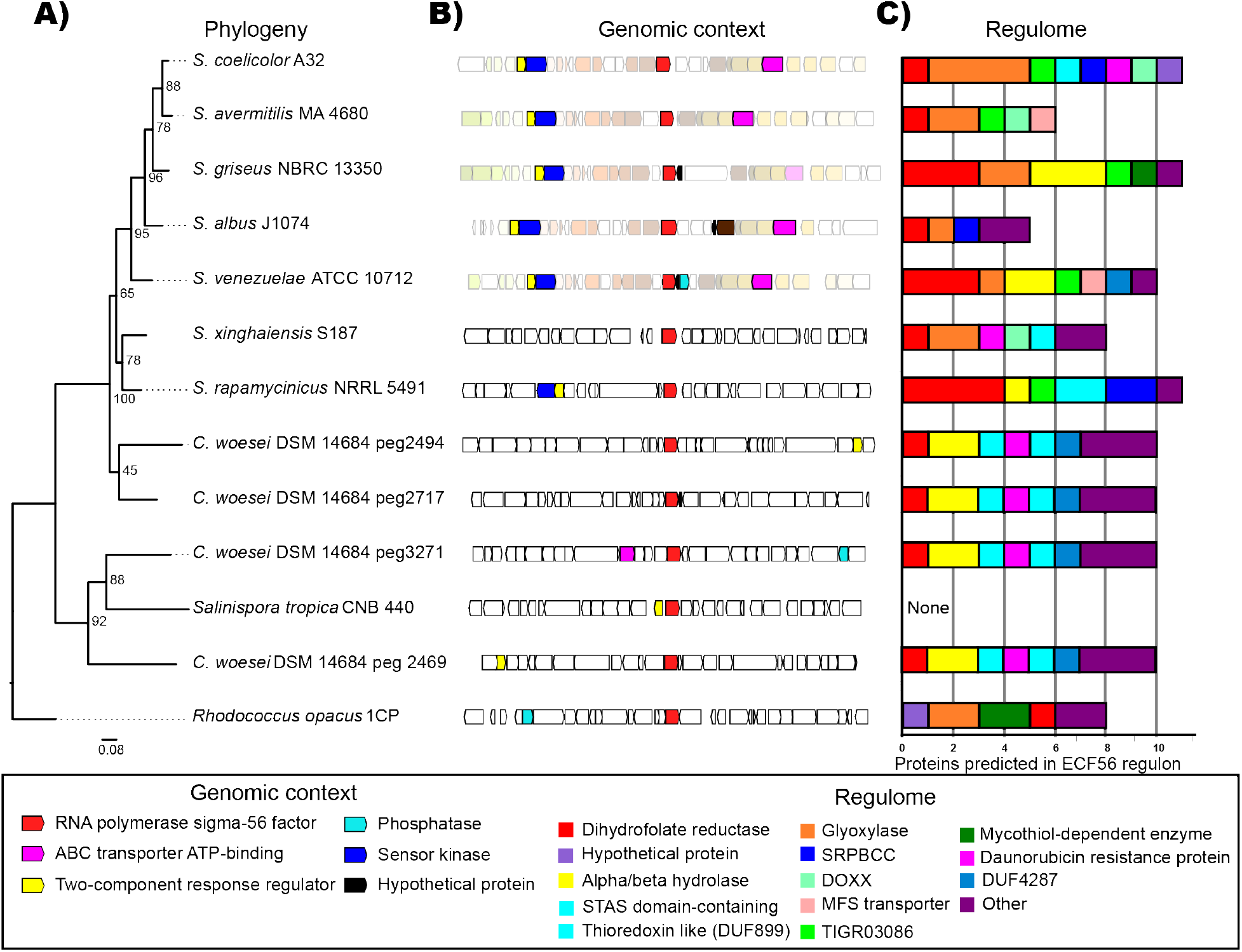
Distribution of ECF56 across Actinobacteria. **A**, Phylogenetic distribution of representative orthologs from selected Actinobacteria species, Most *Streptomyces* genomes include only one copy, while C. woesei includes four. **B**. Genomic neighborhood for each ECF56 ortholog in red. The genes are colored according to their homology. Faded colors represent the genome neighborhood conserved in recently diverged orthologs. Solid colors represent conserved genes within the genomic neighborhood, their annotation is shown at the bottom. **C.** Abundance and functions of the genes regulated by ECF56 in each species. Each bar accounts for the number of the proteins predicted (and confirmed in the case of *S. venezuelae*), to be regulated by ECF56. The functions are represented by different colors shown in the box at the bottom.

**Figure 4.**
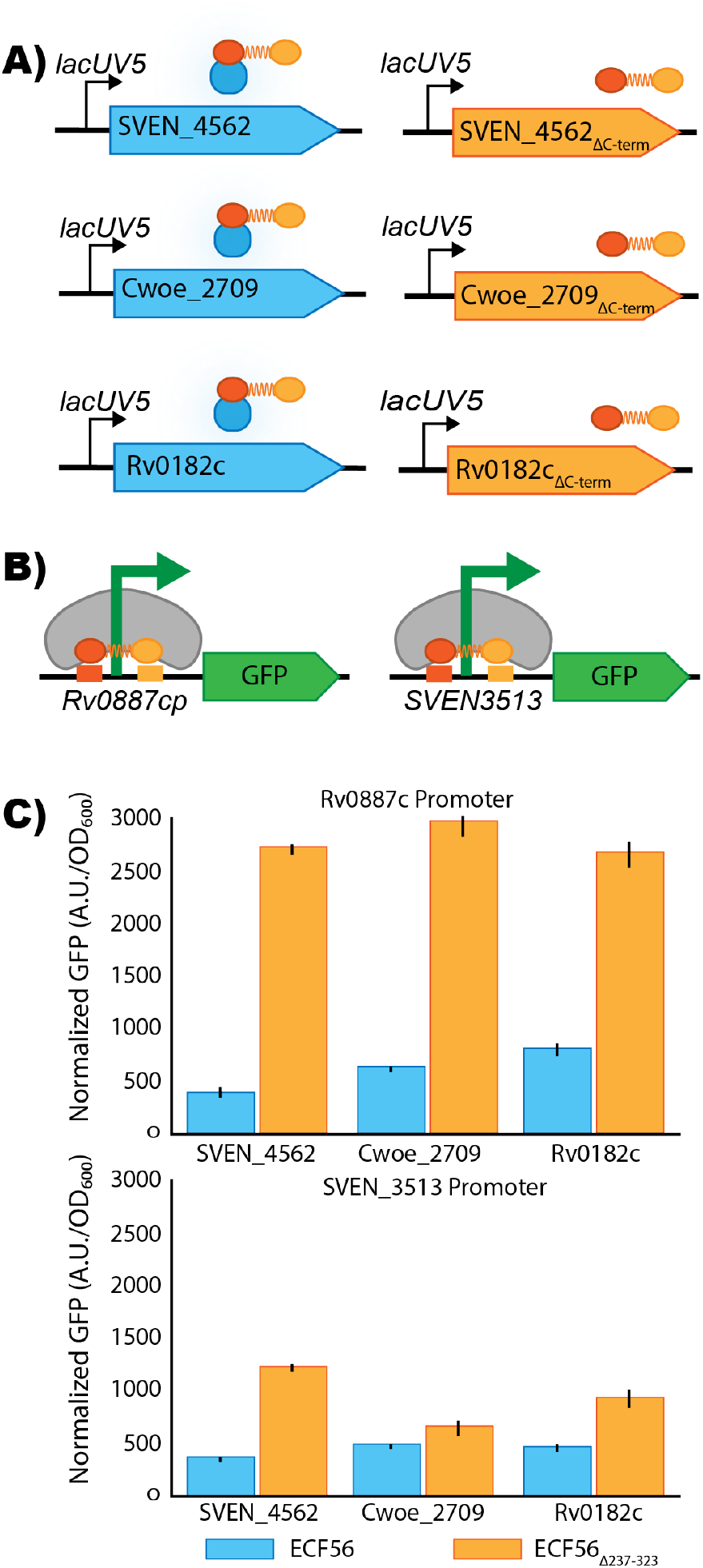
Genetic architecture of ECF56 across phyla. **A)** A wild-type and truncated version of three phylogenetically distinct ECF56 promoters are driven by the lacUV5 promoter were constructed **B)** Promoters from *M. tuberculosis* (RV0887c) and *S. venezuelae* (SVEN_3513) transcribe the GFP reporter in the presence of the ECF sigma factor **C)** *ex situ* assay illustrating the overnight GFP expression levels in *E. coli* normalized to cell density. Error bars represent 95% confidence intervals, n=3.

**Figure 5:**
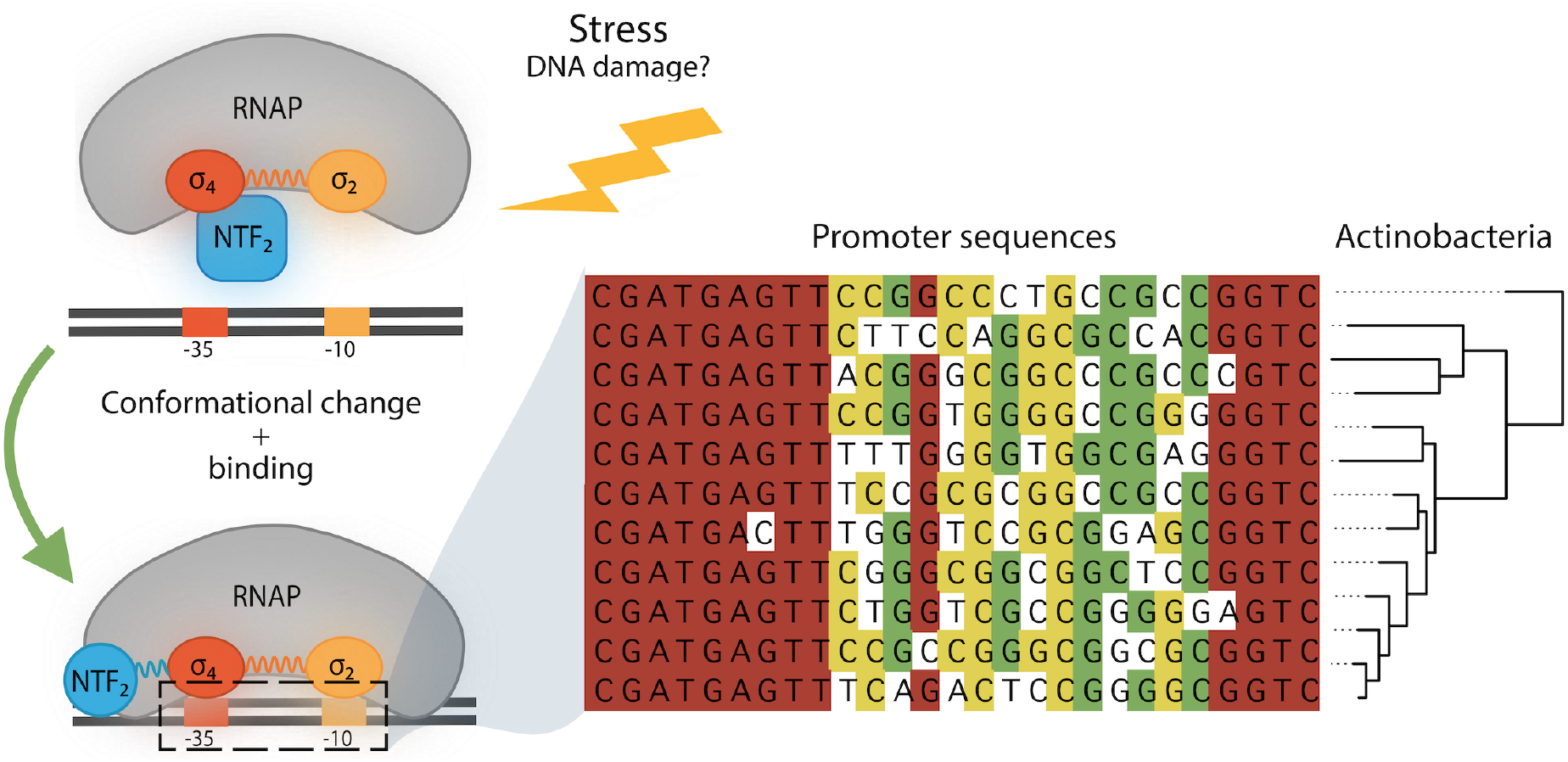
Investigations into sigma factor ECF56 in *S. venezuelae* uncovers that the removal of the inhibitory NTF2 domain results in constitutive transcription of a highly conserved promoter across vast phylogenetic distances.

The regulon of each organism shown in figure 3 was obtained by searching for the ECF56 binding consensus sequence “CGATGAGTTnnnnnnnnnnnnnnnGTC” in each representative genome using the Artemis genome browser. Genes were considered ECF56 regulon members if located immediately downstream of the ECF56 binding sequence. Members of the regulon were manually annotated by performing blast searches against the NR database and searching for annotated domains in the hmmscan server (51).

## Supporting information

Supplemental Attachment 1

Supplemental Attachment 2

## Acknowledgements

The authors would like to thank Matthew Traxler for helpful discussions during the preparation of this manuscript. This work was part of the DOE Joint BioEnergy Institute (https://www.jbei.org) supported by the U.S. Department of Energy, Office of Science, Office of Biological and Environmental Research, and was part of the Agile BioFoundry (http://agilebiofoundry.org) supported by the U.S. Department of Energy, Energy Efficiency and Renewable Energy, Bioenergy Technologies Office, through contract DE-AC02-05CH11231 between Lawrence Berkeley National Laboratory and the U.S. Department of Energy. This work was also funded by National Institutes of Health Awards, F32GM125179. The views and opinions of the authors expressed herein do not necessarily state or reflect those of the United States Government or any agency thereof. Neither the United States Government nor any agency thereof, nor any of their employees, makes any warranty, expressed or implied, or assumes any legal liability or responsibility for the accuracy, completeness, or usefulness of any information, apparatus, product, or process disclosed, or represents that its use would not infringe privately owned rights.

## Contributions

Conceptualization, M.G.T., A.Z.; Methodology, M.G.T., A.Z., P.C.M., T.D.R.;Investigation, M.G.T., A.Z., P.C.M., T.D.R., S.C., R.M.P., G.G., A.N.P., J.F.B., J.M.B., V.R.; Writing – Original Draft, M.G.T, A.Z., P.C.M.; Writing – Review and Editing, All authors.; Resources and supervision N.J.H., P.M.S., J.D.K.

## Supplementary Material

**Figure S1.**
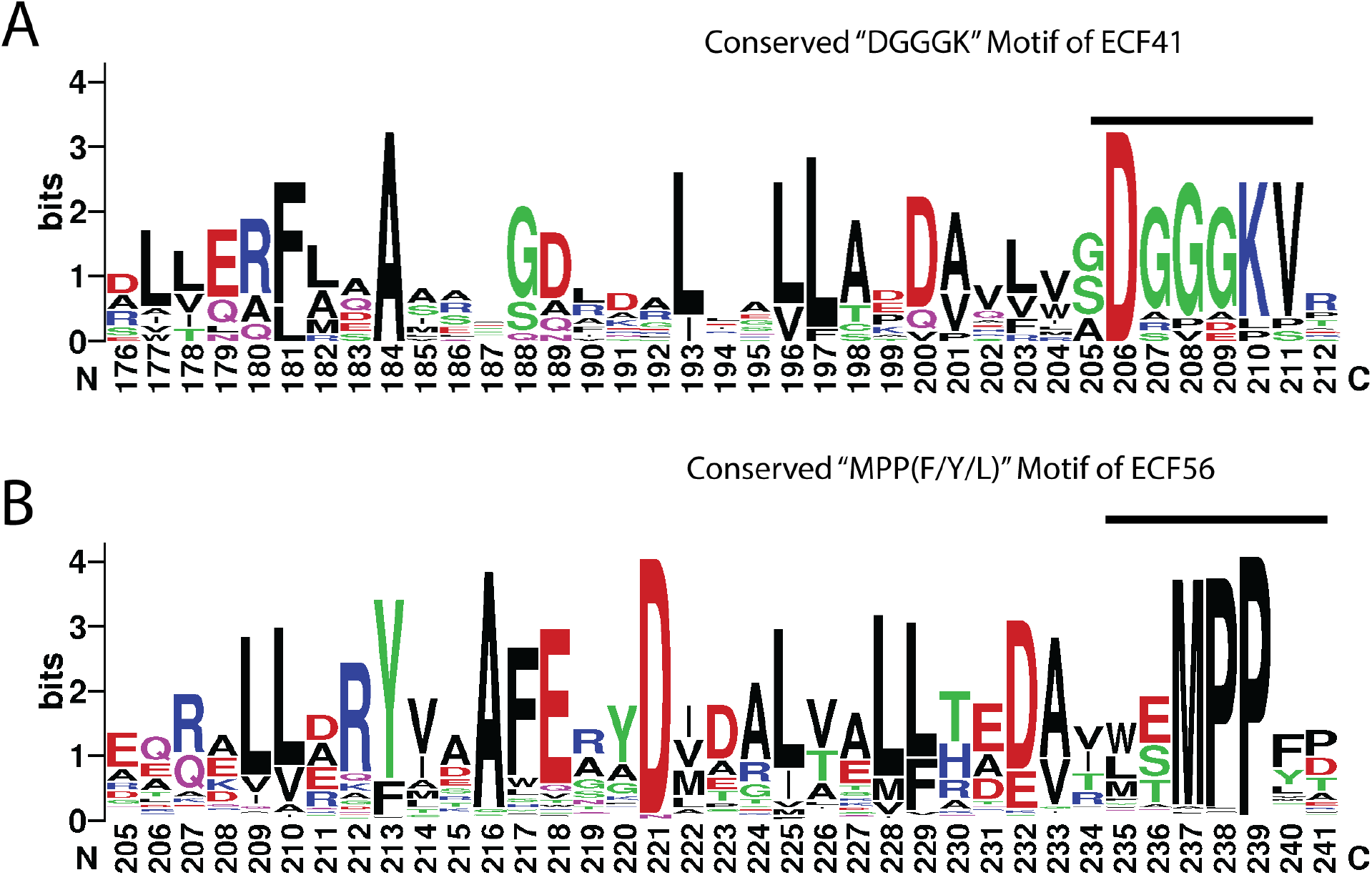
Conserved motifs of ECF41 and ECF56 that delineate truncation sites. Conserved residues of the beginning of the NTF2 domain of both ECF41 (A), and ECF56 (B) sigma factors. The conserved motifs DGGGK of ECF41 and MPP(F/Y/L) of ECF56 are highlighted.

**Table S1:** Suspected read through or overexpression artifacts from RNA-Seq analysis. Genes that were found to be highly upregulated as a consequence of read through during the overexpression of truncated SVEN_4562. The b-value is a measure of change in expression (17), where as the qval is a corrected significance value

| Gene ID | Description | *b* | *q-value* |
| --- | --- | --- | --- |
| SVEN_2281 | 3-oxoacyl-[acyl-carrier protein] reductase | 7.02 | 2.23E-67 |
| SVEN_3512 | Thermophilic serine proteinase precursor | 6.25 | 6.09E-54 |
| SVEN_3511 | hypothetical protein | 6.16 | 6.63E-78 |
| SVEN_6114 | Permease of the drug or metabolite transporter superfamily | 5.44 | 3.63E-71 |
| SVEN_3814 | hypothetical protein | 5.42 | 3.85E-65 |
| SVEN_5314 | putative integral membrane protein | 5.09 | 1.69E-26 |
| SVEN_3510 | Transcriptional regulator, GntR family | 5.04 | 2.09E-70 |
| SVEN_4124 | aminoglycoside phosphotransferase | 4.890 | 4.92E-47 |
| SVEN_3509 | D-3-phosphoglycerate dehydrogenase | 4.47 | 1.69E-62 |
| SVEN_0042 | hypothetical protein | 4.00 | 1.81E-22 |
| SVEN_4562 | RNA polymerase sigma-70 factor | 3.99 | 2.22E-28 |
| SVEN_4122 | secreted protein | 3.61 | 1.32E-46 |

