## Supplementary figures and images for "Characterization of an ECF56-family sigma factor from *Streptomyces venezuelae* reveals a highly conserved regulome"

### Supplemental Attachment 2

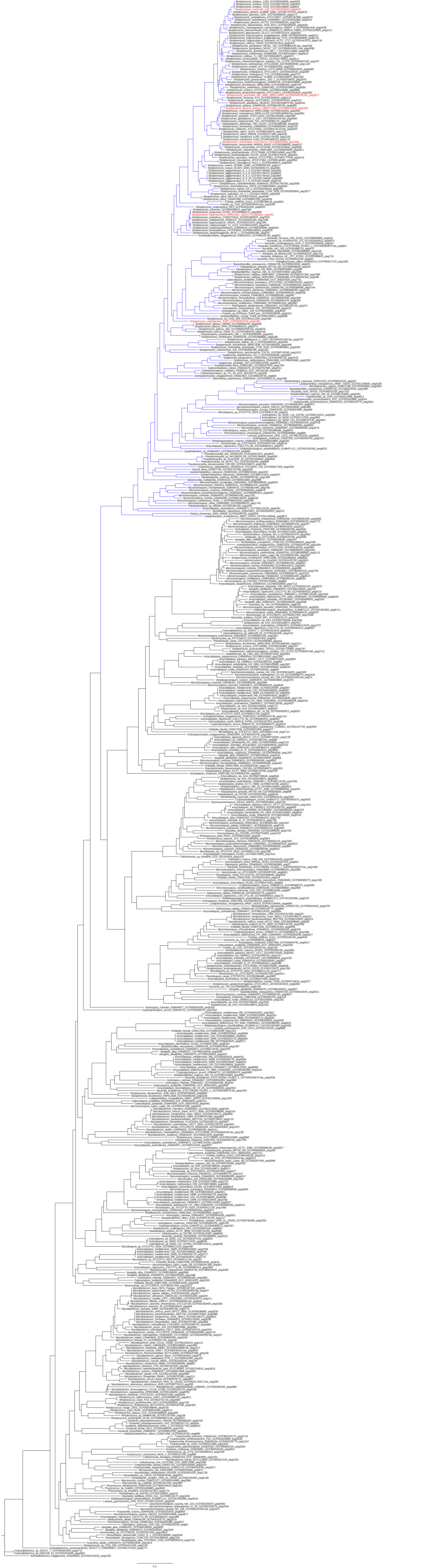
